# Title: “Mitochondrial GWAS and Association of Nuclear - Mitochondrial Epistasis with BMI in T1DM Patients”

**DOI:** 10.1101/436519

**Authors:** Agnieszka H. Ludwig-Słomczyńska, Michał T. Seweryn, Przemysław Kapusta, Ewelina Pitera, Urszula Mantaj, Katarzyna Cyganek, Paweł Gutaj, Łucja Dobrucka, Ewa Wender-Ożegowska, Maciej T. Małecki, Paweł P. Wołkow

## Abstract

Mitochondria are organelles whose main role is energy production and might influence obesity. They are the only organelles with their own genome. Here we have genotyped 435 patients with type 1 diabetes using Illumina Infinium Omni Express Exome-8 v1.4 arrays and performed mitoGWAS on BMI. We have analyzed additive interactions between mitochondrial and nuclear variants in genes known to be associated with mitochondrial functioning (MitoCarta2.0) and confirmed and refined the results on external cohorts - Framingham Heart Study (FHS) and GTEx data. The linear mixed model analysis was performed using the GENESIS package in R/Bioconductor We have found a nominal association between rs28357980 localized to MT-ND2 and BMI (β=−0.69, p=0.056). This was confirmed on 1889 patients from FHS cohort (β =−0.312, p=0.047). Next, we have searched for additive interactions between mitochondrial and nuclear variants. MT-ND2 variants interacted with variants in SIRT3, ATP5B, CYCS, TFB2M and POLRMT genes. TFB2M is a mitochondrial transcription factor and together with TFAM creates transcription promoter complex for mitochondrial polymerase POLRMT. We have found that the interaction between rs3021088 of MT-ND2 gene and rs6701836 in TFB2M has led to BMI decrease (inter_pval=0.0241), while interaction of rs3021088in MT-ND2 and rs41542013 in POLRMT gene led to BMI increase (inter_pval=0.0004). The influence of these interactions on BMI was confirmed on external cohorts. Here, we have shown that variants in mitochondrial genome as well as additive interactions between mitochondrial and nuclear SNPs influence BMI in T1DM and general cohorts.

**Author summary:** Obesity is an epidemic of our times. It is known that it results from an imbalance between energy intake and its expenditure, while mitochondria are organelles whose main role is energy production. They are the only organelles that contain their own genome. Thus, we have genotyped 435 patients with type 1 diabetes and looked on single mitochondrial variant influence as well as on additive interactions between mitochondrial and nuclear variants which might affect BMI. Our analysis has shown, that rs28357980 localized to MT-ND2 is associated with BMI. Next, we looked whether variants in this gene, which builds complex I of the electron transport chain, might interact with nuclear variants and together they modify obesity risk. We focused mainly on mitochondrial biogenesis and found that interactions between variants in TFB2M (rs6701836) or POLRMT (rs41542013) and MT-ND2 (rs3021088) affect patients BMI. TFB2M is a mitochondrial transcription factor which, together with TFAM, creates transcription promoter complex and enables transcription by mitochondrial polymerase POLRMT. The obtained results were also confirmed and refined on external cohorts - Framingham Heart Study (FHS) and GTEx data. Thus, we have shown that variations in mitochondrial genome and its interactions with nuclear variants might have an influence on BMI.

## Introduction

Mitochondria are organelles whose main role is energy production. They are the only organelles that contain their own genome. The mitochondrial genome is a double stranded 16.5 kb long molecule which resembles that of prokaryotes [1]. The two strands differ in nucleotide (G +T) composition - the one which contains more guanines is named heavy (H) strand; the other, which is cystosine-rich is called L-strand (light strand). The mitochondrial genome does not contain any introns and has no spaces between the genes, although it contains ~1000 nucleotide long non-coding control region (displacement loop or D-loop) where origin of replication of the H-strand as well as promoters of transcription, both for H- and L-strands are localized. mtDNA codes for 37 genes. Among them 13 code for polypeptides, the rest - 2 rRNAs (12S and 16S) and 22 tRNAs - are necessary for mitochondrial protein synthesis. All of the 13 mRNAs code for subunits of 4 out of 5 oxidative phosphorylation (OXPHOS) complexes. The rest of peptides which are needed to build electron transport chain (ETC) as well as to maintain mitochondrial functioning is nuclear-encoded [2]. It was shown that mitochondrial function correlates with cells’ metabolic state and can influence obesity [3].

Obesity is an epidemic of our times [4]. Current trends suggest that by the year 2030 more than 50% of Americans will be obese [5], while morbid obesity will affect even 10% of UK population [6]. It was shown that obesity became prevalent even in people that historically are believed to be of rather lean posture (e.g. Type 1 diabetes) [7]. In this cohort overweight/obesity became two times more prevalent over the last 30 years [8,9]. Such a burst in obesity prevalence can, in part, be attributed to lifestyle changes, however obesity is also highly heritable. Twin studies suggest that 40-70% of variability of BMI, which is the most popular measure to assess obesity, can be attributed to genetic variation [10]. Even though large scale genome wide association studies (GWAS) including hundreds of thousands of individuals and millions of autosomal single nucleotide variants have been performed, they led to discovery of only around 100 genetic variants associated with BMI [11–13]. Thus, a substantial part of genetic variability that influences BMI still remains to be discovered, especially that most of the associated variants are non-coding (making explanation of their role in obesity even more difficult).

It is known that obesity results from an imbalance between energy intake and its expenditure. Since it was shown that interaction and communication between nuclear and mitochondrial genomes is indispensable for normal cell function [14,15] it seems reasonable to look for interactions between SNPs in the nucleus and in the mitochondria and their associations with obesity [16–18]. Here, we have performed mitochondrial GWAS as well as analysis of genetic interactions between mitochondrial and nuclear variants which are localized to genes known to have an influence on mitochondrial functioning and we searched for their correlation with BMI.

## Results

### Single SNP analysis – mitoGWAS

We have recruited a group of 435 women who were pregnant or were preparing to conceive for the study on gestational weight gain and its genetic contributors. Here, we have however used the cohort to identify BMI-associated loci in the mitochondrial genome. The mean age of patients was 28.5 years. Most of them were of normal weight and have never been pregnant before being included to the study. Mean disease duration was 12 years, while median insulin daily dose equaled 40 IU. Anthropometric data are presented in Table 1.

**Table 1.**
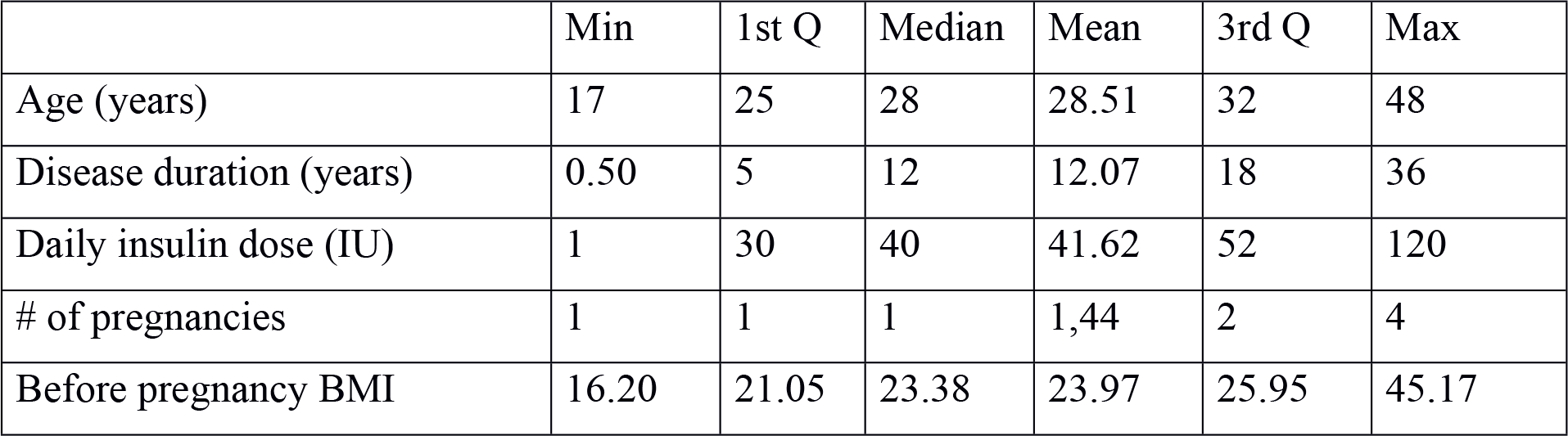
Antropometric data of T1DM cohort.

Forty two mitochondrial variants were included in the analysis (S1a Table). Nominal association with lower BMI was shown for 4 mitochondrial variants. Three of them were localized to non-coding part of mitochondrial genome – one to mitochondrial ribosomal gene MT-RNR2 and two to MT-tRNAs (Arg, Thr). The last one - rs28357980 was localized to MT-ND2 gene (MitoA4917G), which is part of the complex I of electron transport chain. The variant leads to amino acid change from Asparagine to Aspartic acid in position 150 of MT-ND2 protein. Results of mitoGWAS on BMI in T1DM cohort are presented in Table 2.

**Table 2.** mitoGWAS results on BMI in T1DM cohort.

| Probe_ID | rs_number | MAF | Mitochondrial localization | Gene | βSNP | SE | p-value |
| --- | --- | --- | --- | --- | --- | --- | --- |
| exm-rs28358279-132_T_R_1990486714 | rs28358279 | 10.8% | 10463 | MT-TR | -0.73 | 0.355 | 0.040 |
| **exm2263338-0_B_R_1978044975** | **rs28357980** | **10.3%** | **4917** | **MT-ND2** | **-0.69** | **0.362** | **0.056** |
| exm2263337-0_B_R_1978044973 | rs527236198 | 9.9% | 15928 | MT-TT | -0.70 | 0.370 | 0.056 |
| exm2216232-0_T_F_1955482039 | rs2897260 | 9.6% | 1888 | MT-RNR2 | -0.70 | 0.374 | 0.058 |

An analogous analysis of 87 mitochondrial variants (S1b Table) on 1889 subjects gathered in Framingham Heart Study (validation cohort) was performed. The cohort consisted of 1037 women and 852 men. Their mean age was 34.6, half of them were normoglycemic. Median BMI for this cohort was 24.28 (Table 3).

**Table 3.**
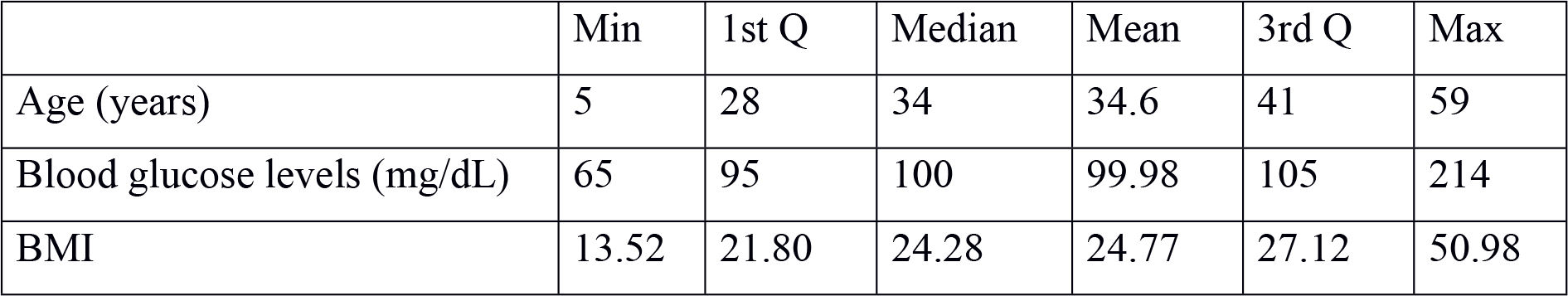
Antropometric data of FHS cohort.

Nominal associations with BMI were found for 7 variants. All nominally associated variants led to BMI decrease. Four of them were coding, the rest was localized to rRNA and tRNAs. Among coding variants we have found the same variant rs28357980 localized to MT-ND2 gene for which an association with BMI in T1DM cohort was found. Results of mitoGWAS on BMI in FHS cohort are presented in Table 4.

**Table 4.** mitoGWAS results on BMI on FHS cohort.

| Probe_ID | rs_number | MAF | Mitochondrial localization | Gene | βSNP | SE | p-value |
| --- | --- | --- | --- | --- | --- | --- | --- |
| MitoG3012A | rs3928306 | 45.8% | 3010 | MT-RNR2 | -0.241 | 0,113 | 0.033 |
| **MitoA4918G** | **rs28357980** | **19.1%** | **4917** | **NT-ND2** | **-0.312** | **0.157** | **0.047** |
| MitoT9900C | rs41345446 | 3.8% | 9899 | MT-CO3 | -0.785 | 0.338 | 0.020 |
| MitoA11252G | rs869096886 | 35.8% | 11251 | MT-ND4 | -0.241 | 0.123 | 0.051 |
| 200610-74 | rs193302994 | 36.1% | 15452 | MT-CYB | -0.242 | 0.123 | 0.049 |
| MitoG15929A | rs527236198 | 18.2% | 15928 | MT-TT | -0.396 | 0.163 | 0.015 |
| MitoA16164G | rs41479950 | 4.6% | 16163 | MT-TP | -0.689 | 0.310 | 0.027 |

### MT-ND2 gene interactions within MitoCarta

Since each of the analysis we performed (mitoGWAS in both cohorts) directed us towards MT-ND2 gene, we have looked closer into its interactions with genes listed in MitoCarta (S2 Table). Most significant interactions for MT-ND2 were associated with TCA cycle and respiratory electron transport, metabolism of amino acids and mitochondrial biogenesis (S3 Table). The last category can potentially have the most crucial influence on mitochondrial functioning and can have profound significance for energy metabolism. Reactome lists 17 genes (POLRMT, TFB1M, TFAM, MTERF, PERM1, TFB2M, ATP5B, SSBP1, SIRT3, POLG2, PPARGC1A, CYCS, NRF1, ALAS1, PEO1, GABPA, ESRRA) which constitute this pathway, five of which are present in the results of our analysis. The list of MT-ND2 variants interactions within this pathway is presented in Table 5.

**Table 5.** List of MT-ND2 interactions within mitochondrial biogenesis pathway.

| Probe_ID | rs_number | MitoMAF | Nuclear SNP ID | Nuclear MAF | Nuclear localization | p-value |
| --- | --- | --- | --- | --- | --- | --- |
| exm2263338-0_B_R_1978044975 | rs28357980 | 10% | rs113919457 | 19.3% | SIRT3 | 0.0016 |
|  |  |  | rs11607019 | 19.3% | SIRT3 | 0.0016 |
| exm-rs3021088-132_B_R_1990477615 | rs3021088 | 5.1% | rs2950393 | 28.4% | ATP5B | 0.0028 |
|  |  |  | rs10215217 | 28.9% | CYCS | 0.001 |
|  |  |  | rs41542013 | 15.0% | POLRMT | 0.0004 |
|  |  |  | rs6701836 | 37.0% | TFB2M | 0.024 |

We looked closer into these interactions and found that variant rs3021088 of MT-ND2 gene interacted with variants in TFB2M (rs6701836) and POLMRT (rs41542013) genes, both of which are necessary for mitochondrial transcription. TFB2M is a mitochondrial transcription factor which, together with TFAM, creates transcription promoter complex and enables transcription by mitochondrial polymerase POLRMT (S1 Figure).

When looking at the interaction between TFB2M nuclear variant - rs6701836 and MT-ND2 mitochondrial variant - rs3021088 we found that the combination of the two led to BMI decrease [(nuc_eff(single model)=-0.0660; nuc_pval(single)=0.857, joint_eff(inter)=0.0874, nuc_pval(inter)=0.0773, inter_eff=-2.225, inter_pval=0.0241]. Neither nuclear nor mitochondrial variant on its own had such an effect [p(nuc) - p=0.8577, p(mito) – p=0.116].

Interaction between POLRMT variant rs41542013 and MT-ND2 mitochondrial variant rs3021088 led to BMI increase [nuc_eff(single model)=0.296; nuc_pval(single)=0.546, nuc_eff(inter)=-0.085, joint_pval(inter)=0.0016, inter_eff=4.015, inter_pval=0.0004]. None of these variants on its own was not associated with BMI.

Thus, we have found an example of negative interaction between MT-ND2 and TFB2M and positive interaction between MT-ND2 and POLRMT variants.

### Functional significance

Next, we assessed the functional significance of these interactions. Mitochondrial rs3021088 is a missense variant which leads to Alanine to Threonine substitution in position 331 of MT-ND2 protein.

eQTL data (GTEx) for nuclear rs6701836 showed that it influenced TFB2M expression in thyroid gland. POLRMT variant rs41542013 in subcutaneous adipose tissue leads to lower POLRMT expression (Figure 1). In GTEx data, mRNA levels of MT-ND2 and TFB2M correlated with higher BMI [p(nuc) = 0,0269 eff_nuc=2.465e-01, p(mito)= 0,0244, eff_mito=2.243e-04] in liver tissue, however, their interaction led to decrease of BMI [p=0,0308, inter_eff=-1.009e-05]. In GTEx data, mRNA levels of MT-ND2 and POLRMT on its own correlated with lower BMI [p(nuc) = 0.0492 eff_nuc=-2.386e-01, p(mito)= 0.0688, eff_mito=-2.233e-04] in subcutaneous adipose tissue, however, their interaction led to increase of BMI [p=0.0235, inter_eff=1.023e-05]. Taken together, the interactions on the mRNA level are in line with what was discovered in the T1DM cohort.

**Figure 1.**
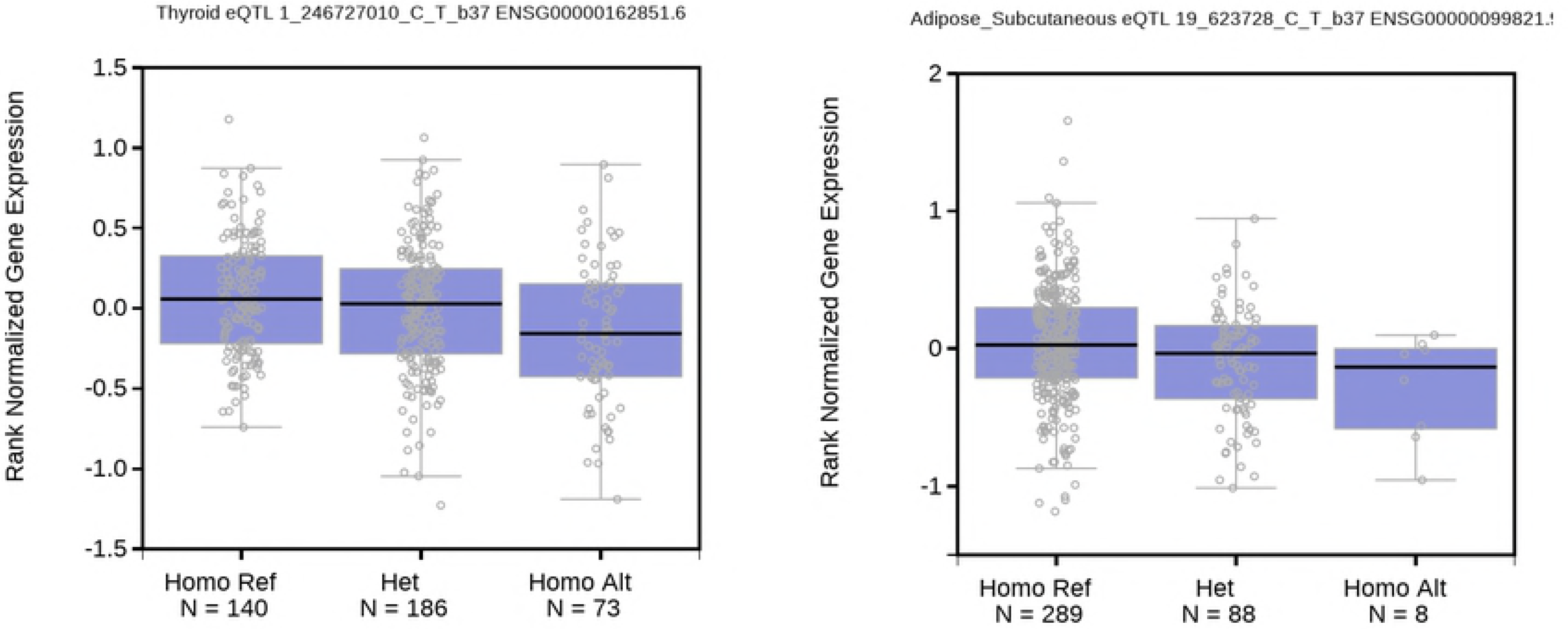
Influence of rs6701836 on expression of TFB2M gene and rs41542013 on POLRMT expression.

### Validation cohort

We also used FHS as a validation cohort to confirm the epistatic interactions between SIRT3, ATP5B, CYCS, TFB2M or POLRMT and MT-ND2.

We were able to confirm the interactions and their influence on BMI for SIRT3 and ATP5B genes. For SIRT3 we have found an interaction between mitochondrial variant localized to 5460 position (MAF 10.1%) and two nuclear variants - rs11602954 (p=0.03) and rs11606393 (p=0.028). These variants are in LD with each other and with previously described variants in SIRT3 (in Table 5). For ATP5B gene we have found that mitochondrial variant localized to position 4769 interacted with seven variants (rs2255074, rs1107479, rs7973157, rs2950390, rs2958154, rs2290893, rs2035081) which were in LD with rs2950393.

We looked closer into interactions of MT-ND2 and TFB2M or POLRMT. For TFB2M gene, we have found two interactions. Variant rs10924779 interacted with MT-ND2 variant localized to 5460 position (MAF=10.1%). Their interaction leads to BMI increase (p=0.037), while each of these variants on their own did not have an influence on BMI. Variant rs4654291 interacts with the same mitochondrial variant and leads to BMI increase (p=0.045), while each of these on its own does not affect BMI. For POLRMT gene we have identified an intronic variant rs10411491 (MAF = 3%) which interacted with mitochondrial variant positioned at 4647 (MAF=4.8%). Their interaction led to BMI increase, although due to low intronic variant MAF it did not reach statistical significance.

Thus, we can conclude that interactions between TFB2M or POLRMT and MT-ND2 gene might affect BMI.

### Whole genome MT-ND2 interactions

Next, we looked into whole genome MT-ND2 interactions. All variants which entered statistically significant interactions with MT-ND2 gene in FHS cohort and T1DM cohort were listed (S4a,b Table). These variants were compared to results gathered in GWAS catalog. Among 366 nuclear variants which interacted with mitochondrial variant rs28357980, 5 were found in GWAS Catalog and 2 of them met genome wide significance threshold. Out of 1157 variants which interacted with mitochondrial variant rs3021088, 35 were listed in GWAS Catalog and 13 were significant at the level of p<e10-09 (S4c Table). According to GWAS Catalog these variants were associated, among others, with blood metabolite levels, blood protein levels, LDL and total cholesterol, bone mineral density, age-related hearing impairment.

## Discussion

Functioning of mitochondria have a profound effect on whole organism. The variants within mitochondrial as well as nuclear genomes might have an influence on the levels of ATP produced within mitochondria. ATP depletion can induce endoplasmatic reticulum (ER) stress and lead to reactive oxygen species (ROS) generation, which later on will result in mitochondrial DNA damage and create a vicious cycle of mitochondrial inefficacy. Here we have looked for mitochondrial variants and their interaction with nuclear variants which may be associated with BMI and diabetes.

Our mitoGWAS analyses have shown that both protein coding and non-coding variants are associated with BMI. We have however looked closer into the former, as they might directly influence electron transport chain (ETC) and cells energy production. The analysis of T1DM cohort led to discovery of a variant in MT-ND2 gene. The predictive algorithms do not suggest that rs28357980 (G4917A) might be damaging (SIFT - 0.09, PolyPhen - 0.06), however the variant has been shown to be associated with multiple sclerosis [19] and colorectal cancer [20]. The G4917A variant is one of the defining variants of haplogroup T, which in 2013 was shown to be a risk factor for morbid obesity [21]. Moreover the variant is non-synonymous and affects the protein by substitution of highly conserved Aspartate at amino acid 150 to an Asparagine. What is more interesting, our analysis of the validation cohort (FHS) confirmed the association of the same variant with BMI.

rs28357980 (G4917A) is localized to MT-ND2 gene which is part of complex I of ETC. It is the first site of oxidative phosphorylation. Complex I is built of 46 proteins, 7 of which are mitochondrially coded (ND1, ND2, ND3, ND4, ND4L, ND5, ND6) and form very hydrophobic subunits within mitochondrial membrane. Disruptive mutations in ND subunits are commonly found as somatic mutations in tumors, but are not found as germline mutations associated with human diseases, due to their lethality [22]. Previous studies have already associated variants in these genes with obesity. A paper from 2014 has shown, apart from associations with two other positions, an association between three variants in complex I genes– MT-ND1, MT-ND2 and MT-ND4L [23]. Moreover, the variant C5178A in MT-ND2 gene was shown to lead to lower incidence of autoimmune diabetes. The A allele was protective against both autoimmune and alloxan-induced free radical-mediated diabetes in mice, possibly by suppressing ROS production at the β-cell level [24–26]. Apart from metabolic disorders, mutations in genes of complex I of ETC were shown to be associated with childhood acute lymphoblastic leukemia or were shown to be a poor prognostic factor in oral cancer [27].

Since several GWAS data show that a substantial part of genetic variability of obesity is still unknown one might suspect that it is hidden in more complex associations, meaning genetic interactions. It is known that mitochondrial functioning is a result of anterograde and retrograde signaling between mitochondrial and nuclear genomes. Most of the studies performed by now, did not analyze point mitochondrial variants, but conplastic animals in which nucleus from one organisms was fused with cytoplasts of the other. Such experiments have shown that introduction of exogenous mitochondria (in which mitochondrial genome differed from the original by one or few nucleotides) influenced organismal phenotype. For example, conplastic rats where shown to have impaired glucose tolerance, while mice were more resistant to experimental autoimmune encephalomyelitis [28], had disrupted activity of the components of TCA cycle [29] or altered mitochondrial and cellular adaptation during aging [30]. Moreover, the mutation in MT-ND2 gene (C4738A) in mouse fibroblasts led to significantly higher mitochondrial complex I activity, enhanced ATP production, reduced ROS production with similar MT-ND2 protein expression levels [31]. A lot of studies have shown that even a point mutation in mitochondrial genome which was introduced onto another nuclear background led to severe mitochondrial dysfunction. A point mutation in ATP8 gene (7778 G/T) in C57BL/6N-mt^FVB/N^ mice led to lower insulin secretion in isolated islets after glucose stimulation when compared with C57BL/6N-mt^AKR/J^ mice. It also had reduced mitochondrial function in brain, spleen and liver [32] as well as showed a 3-fold higher generation of mitochondrial ROS production compared to C57BL/6N-mt^AKR/J^ mice [33].

However, since mitochondrial genome mutates faster than nuclear (for example due to ROS proximity) the incompatibility between the two genomes might occur during organismal lifetime. What is more, these interactions can also be influenced by the environment. Thus, we have looked into epistatic interactions of MT-ND2 variants in our T1DM cohort. The Reactome analysis has shown an enrichment of interactions that are associated with mitochondrial biogenesis. Our data show that POLRMT and TFB2M variants interact with variants in MT-ND2 and affect BMI. POLRMT and TFB2M are genes that act in concert to perform mitochondrial transcription and replication thus variation within their sequence, and simultaneously in the mitochondrial DNA sequence, can have an influence on mitochondrial and gene copy number as well as influence efficacy of the two processes [34–36]. Since it is very difficult to assess the significance of such interactions we also looked into all interactions of MT-ND2 variants and checked whether nuclear variants which interacted with mitochondrial MT-ND2 gene were listed as significant in any GWAS study performed until today, as we believed this would strengthen our findings. Our analysis has confirmed that some of the nuclear variants were significantly associated with traits which are part of obesity phenotype, e.g. cholesterol level, blood metabolites level or with diseases which are known to be influenced by mitochondrial deficiencies e.g. hearing loss.

In conclusion, here we find that rs28357980 localized to MT-ND2 gene of mitochondrial genome is associated with BMI both in T1DM and in general cohort. What is more, we show that genetic epistasis might influence obesity phenotype by interaction of variants in MT-ND2 gene with nuclear variants in genes responsible for mitochondrial replication and transcription.

## Methods

### Patients

Patients were recruited either in Department of Metabolic Diseases University Hospital in Krakow or in Division of Reproduction Department of Obstetrics, Gynecology and Gynecological Oncology, Poznan University of Medical Sciences. All patients enrolled to the study were young women with type 1 diabetes (T1D) and on insulin treatment. Whole blood samples were drawn and stored at −80ºC. This study was approved by the Bioethical Committees of the Jagiellonian University and Poznan University of Medical Sciences and performed according to the Helsinki Declaration. Written informed consent was collected from all patients.

### Genotyping

DNA was extracted from whole blood with the use of automated nucleic acid extraction system Maxwell (Promega). Five hundred twenty seven samples were genotyped on Illumina Infinium Omni Express Exome-8 v1.4 arrays according to manufacturer’s instructions.

### Quality Control (QC)

Genotypes were called by GenomeStudio software Genotyping module (version 2.0, Illumina Inc.) according to manufacturers’ instructions (Technical Note: Genotyping, Infinium ^®^ Genotyping Data Analysis, www.illumina.com) Briefly, we removed all samples with 10%GenCall score <0.4, Call Rate <0.95 and discordant sex information. Then, all SNPs on chrX, chrM, chr0 were removed and hard cut off metrics was applied. Next, we manually curated 201 variants on chrM also in Genome Studio, recalculated statistics and excluded samples with CallRate <0.99 (180 remained in the analysis). Remaining nuclear genotypes for 940911 SNPs and 513 samples were exported as custom report using plink (version 1.09) [37].

Second step of quality check was performed using custom rscript in Rstudio (version 1.1.383) [38] with R (version 3.4.2) [39] according to plink manufacturers’ instructions and KING [40]. Briefly, we removed duplicates with lower call rate and samples with cryptic kinship, equivalent to third-degree relatives and higher (kinship coefficient >0.0442). In addition, we also checked for heterozygosity rate of autosomal SNPs and removed variants with deviation from the Hardy-Weinberg equilibrium (HWE) with P< 1e10-5. The last part of QC was based on EIGENSOFT [41,42] in order to determine population structure in our samples based on HapMap3 dataset [43]. Samples that failed to qualify as European population (CEU) were removed. Finally 476 individuals with 940374 nuclear variants passed QC and were included in the analysis. Apart from nuclear variants, 180 mitochondrial variants were included in the analysis.

### Imputation

The QC-filtered genotype data were checked against reference panel of the Haplotype Reference Consortium (HRC r1.1 2016) [44] with “HRC/1KG Imputation Preparation and Checking Tool” (v.4.2.9.) [45] to exclude strand coding issues during the imputation step. Imputation of QC-filtered genotypes was performed on Michigan Imputation Server (using Minimac3) [46] with the HRC reference panel. Phasing was performed with ShapeIT (version 2.r790) [47]. The Minimac3 output dosage files were then converted to hard calls in PLINK. For further analysis imputed MitoExome variants were chosen based on MitoCarta2.0 gene localization [48].

### Framingham Heart Study Cohort

Genotypes, and clinical phenotypes were acquired from the Framingham Heart Study (FHS) via dbGaP Project #5358 (dbGaP accession number phs000007). Genotypes were taken from the SHARe substudy that used the OMNI5M genotyping array. We considered age (phv00177930), gender (phv00177929), blood glucose (phv00007558) as covariates in our models. Our access of this study was approved by the Ohio State University IRB (Protocol #2013H0096).

### GTEx

Tissue specific RNA sequencing and phenotype data was acquired from the Genotype and Tissue Expression Project (GTEx) via dbGaP Project #5358 (dbGaP accession number phs000424). RNA sequencing had been generated using poly-adenylated priming with reads aligned to HG19, Gencodev12. For further details see Lonsdale et al. [49] and the GTEx website (http://www.gtexportal.org/home/documentationPage). We considered subcutaneous adipose and thyroid gland. Testing for eQTLs was performed via the GTEx online search tool [50].

### Statistical analysis

The linear mixed model analysis was performed using the GENESIS package in R/Bioconductor [51]. In brief, first the genetic relatedness matrix was estimated via the PC-AiR and PC-Relate methods, subsequently a linear model was build and Wald’s test was used to assign significance. The interactions between variants was modeled via the option ‘ivars’ in function ‘assocTestMM’.

## Acknowledgements

The Genotype-Tissue Expression (GTEx) Project was supported by the Common Fund of the Office of the Director of the National Institutes of Health. Additional funds were provided by the NCI, NHGRI, NHLBI, NIDA, NIMH, and NINDS. Donors were enrolled at Biospecimen Source Sites funded by NCI\SAIC-Frederick, Inc. (SAIC-F) subcontracts to the National Disease Research Interchange (10XS170), Roswell Park Cancer Institute (10XS171), and Science Care, Inc. (X10S172). The Laboratory, Data Analysis, and Coordinating Center (LDACC) was funded through a contract (HHSN268201000029C) to The Broad Institute, Inc. Biorepository operations were funded through an SAIC-F subcontract to Van Andel Institute (10ST1035). Additional data repository and project management were provided by SAIC-F (HHSN261200800001E). The Brain Bank was supported by a supplements to University of Miami grants DA006227 & DA033684 and to contract N01MH000028. Statistical Methods development grants were made to the University of Geneva (MH090941 & MH101814), the University of Chicago (MH090951, MH090937, MH101820, MH101825), the University of North Carolina - Chapel Hill (MH090936 & MH101819), Harvard University (MH090948), Stanford University (MH101782), Washington University St Louis (MH101810), and the University of Pennsylvania (MH101822).

The Framingham Heart Study is conducted and supported by the National Heart, Lung, and Blood Institute (NHLBI) in collaboration with Boston University (Contract No. N01-HC-25195). This manuscript was not prepared in collaboration with investigators of the Framingham Heart Study and does not necessarily reflect the opinions or views of the Framingham Heart Study, Boston University, or NHLBI. Additional funding for SABRe was provided by Division of Intramural Research, NHLBI, and Center for Population Studies, NHLBI. Funding for SHARe Affymetrix genotyping was provided by NHLBI Contract N02-HL-64278. SHARe Illumina genotyping was provided under an agreement between Illumina and Boston University. The bioinformatic analysis was performed using Prometheus (AGH, Krakow, Poland), Michigan Imputaton Server (Michigan, MI, USA) and Ohio Supercomputer Center (Columbus, OH, USA).

## Supporting Information Legends

**Table S1a.** List of mitochondrial variants included in the analysis in T1DM cohort

**Table S1b.** List of mitochondrial variants included in the analysis in FHS cohort

**Table S2a.** All interactions of MT-ND2 gene with MitoCarta genes in T1DM cohort

**Table S2b.** Interactions of MT-ND2 gene significant after all corrections of MT-ND2 variant rs28357980

**Table S3.** List of most significant associations of MT-ND2 gene

**Table S4a.** All interactions of MT-ND2 gene genome wide in T1DM cohort

**Table S4b.** All interactions of MT-ND2 gene genome wide in FHS cohort

**Table S4c**. Significant associations of MT-ND2 variants in GWAS Catalog

**Figure S1** - MitoGWAS on BMI type 1 diabetes patients and additive interactions between mitochondrial and nuclear variants in T1DM patients and FHS cohort.

Author contributions
UM, KC, PG and ŁD gathered study cohort and phenotypic data; EWO and MTM supervised cohort construction; AHLS and EP performed genotyping; AHLS, MTS and PK analyzed and interpreted data; AHLS, MTS, PK and PPW wrote the paper; AHLS and PPW and guarantors of this work. All authors contributed to critical revision of the manuscript and approved its publication.

